# Msc1 accumulates at and excludes nuclear pore complexes from nucleus-vacuole junctions

**DOI:** 10.64898/2026.06.03.729915

**Authors:** Sara Medina-Suárez, Héctor M. Estévez-Silva, Noelia Rodríguez-Herrera, Félix Machín

## Abstract

Msc1 is a yeast nuclear envelope (NE) protein that facilitates DNA double-strand break repair. In its absence, cells exhibit abnormal nuclear morphologies and maldistribution of nuclear pore complexes (NPCs). Msc1 is not uniformly distributed across the NE but concentrates onto dynamic patches that often coincide with blebs or herniations. Here, we report that Msc1 abundance and the number of patches dramatically increase after the diauxic shift. Msc1 patches are devoid of NPCs and fully colocalize with nucleus-vacuole junctions (NVJs), which are involved in piecemeal micronucleophagy. In the absence of Msc1, abnormal NPC aggregates accumulate adjacent to vacuoles, both at and outside the NE. We conclude that Msc1 is the key factor that maintains NPC homeostasis as cells prepare to enter quiescence.

## Introduction

Quiescence is an evolutionarily conserved resting state in which proliferative activity is suspended and cellular physiology is largely remodeled (Marescal and Cheeseman, 2020; Sagot and Laporte, 2019). In multicellular organisms, particularly mammals, quiescence is central to tissue homeostasis and stem-cell preservation, and it is now recognized as a regulated metabolic and transcriptional program rather than a passive absence of growth. In microbes, including the budding yeast, a closely related physiological state emerges as cultures exhaust nutrients and progress out of exponential growth into stationary phase (Breeden and Tsukiyama, 2022; Gray et al., 2004; Sagot and Laporte, 2019; Werner-Washburne et al., 1993). Although the stationary phase in yeast is not identical to mammalian quiescence, the two states reflect a shift from growth-centered metabolism to a maintenance-oriented program that supports survival under unfavorable conditions. This conceptual overlap has made yeast a valuable model for dissecting the principles that govern quiescent states across eukaryotes.

In budding yeast, growth begins with a lag phase during which cells adapt to the new environment and reconfigure metabolism before net proliferation becomes evident. Cells then enter logarithmic growth (log phase), in which division occurs at a near-constant rate and population size increases exponentially. As nutrients and other resources progressively decline, cultures pass through a deceleration phase in which growth slows and stress-adaptive programs become increasingly prominent. Stationary phase proper is reached when division and death are approximately balanced and total cell density stabilizes. In between, and when yeast grow under fermentable carbon sources such as glucose, a diauxic shift occurs and marks the metabolic transition when glucose becomes limiting and cells switch from fermentative growth to respiratory utilization of ethanol and other non-fermentable by-products of glycolysis (Breeden and Tsukiyama, 2022; Herman, 2002). During the ensuing post-diauxic growth phase, the cells still grow and divide, though more slowly, until these alternative carbon sources are finally exhausted.

Among the stress-adaptive programs that cells activate upon entry into quiescence, autophagy is essential for maintaining long-term survival. Thus, autophagy supports recycling of macromolecules and organelles to preserve viability during starvation-like conditions (Abeliovich and Klionsky, 2001). In yeast, both bulk autophagy and selective autophagy pathways are activated as resources decline. One of the selective pathways is piecemeal microautophagy of the nucleus (PMN), in which small portions of the nucleus are pinched off and degraded inside the vacuole. PMN occurs at nucleus-vacuole junctions (NVJ) formed by proteins such as Nvj1 on the nuclear envelope and Vac8 on the vacuolar membrane (Farré et al., 2009; Pan et al., 2000; Roberts et al., 2003). PMN removes nonessential nuclear material, including parts of the nuclear envelope (NE) and portions of the nucleolus, which the vacuole then digests (Enkhbaatar et al., 2023). When NVJs are formed, the vacuole membrane invaginates to engulf a teardrop-like bleb from the nucleus. During PMN, nuclear pore complexes (NPCs) are generally excluded from the NVJs (Lee et al., 2020; Pan et al., 2000; Tomioka et al., 2020). For instance, upon treatment with rapamycin, a TORC1 inhibitor that force cells to activate multiple autophagic pathways (including PMN), NPCs were noted to be excluded from the NVJs, although deleting Nvj1 did not block NPC degradation, which argues that NPC turnover occurs through PMN-independent mechanisms such as receptor-mediated nucleophagy or NPC-phagy (Tomioka et al., 2020). At present, there is no known mechanism for NPC exclusion from NVJs. The best-supported idea is that NPCs are kept out of PMN sites because NVJ themselves exclude them. In other words, NPCs are not known to be protected by a dedicated shielding factor.

Msc1 is a NE protein that we have recently identified as a critical factor for maintaining nuclear homeostasis during DNA double-strand break repair (Medina-Suárez and Machín, 2025; Medina-Suárez et al., 2024). In the absence of Msc1, cells exhibit abnormal nuclear morphologies, characterized by excessive blebbing, lobulation and over-compartmentalization of nuclear DNA (Medina-Suárez and Machín, 2025). In accordance with these phenotypes, Msc1 is not uniformly distributed in a subset of growing cells but concentrates into highly dynamic patches that often coincide with blebs or herniations. Another defining feature of Msc1 absence is the misdistribution of NPCs, which tend to aggregate into clusters resembling the Storage of Improperly assembled NPCs (SINC) (Medina-Suárez and Machín, 2025). SINCs are NE-associated compartments that sequester misassembled nucleoporins (Nup proteins) to prevent their incorporation into functional NPCs (Webster et al., 2014). Msc1 has two orthologs in *Schizosaccharomyces pombe*, Ish1 and Les1. In accordance with an expected role in NPC regulation, Les1 has been shown to concentrate onto stalks devoid of NPCs during karyokinesis (Dey et al., 2020). Msc1, Les1, and Ish1 all expose their active globular domains to the NE lumen (Asakawa et al., 2022; Dey et al., 2020; Medina-Suárez and Machín, 2025).

Here, we demonstrate that Msc1 patches markedly increase as cells progress into the stationary phase. We also show that these patches perfectly colocalize with NVJs and are devoid of NPCs. Upon Msc1 depletion, NPCs aggregates both at and out of the NE, are no longer excluded from NE-VM contact sites, and appear to invaginate into the vacuole. We conclude that Msc1 contributes to NPC homeostasis as cells progress into quiescence.

## Results and Discussion

### Msc1 is enriched at nucleus-vacuole junctions after the diauxic shift

In a previous work, we found that Msc1 is a NE protein whose abundance and distribution strongly varies between cells gathered from an asynchronous culture in the log phase (Medina-Suárez et al., 2024). Thus, Msc1-YFP (Msc1 tagged at C-terminal with YFP) levels are often too low to be detected, although it can become more visible in cells synchronized in late mitosis. In the latter, Msc1 could appear either evenly distributed across the NE or concentrated into one or more NE subdomains (patches) (Medina-Suárez and Machín, 2025; Medina-Suárez et al., 2024). In contrast to the log phase (24 h, day 1), Msc1 levels rise sharply in the NE afterwards (Figure 1A, B), and Msc1 progressively concentrates into discrete patches (Figure 1A, C). Thus, there are approximately 50% of nuclei with patches by 48 h (day 2), whereas this fraction rises to ∼70% by 72 h (day 3), and ∼90% at 96 h (day 4). Most nuclei contain just one patch, yet two or more can also be present in a significant proportion (up to 30% of nuclei by day 4).

**Figure 1.**
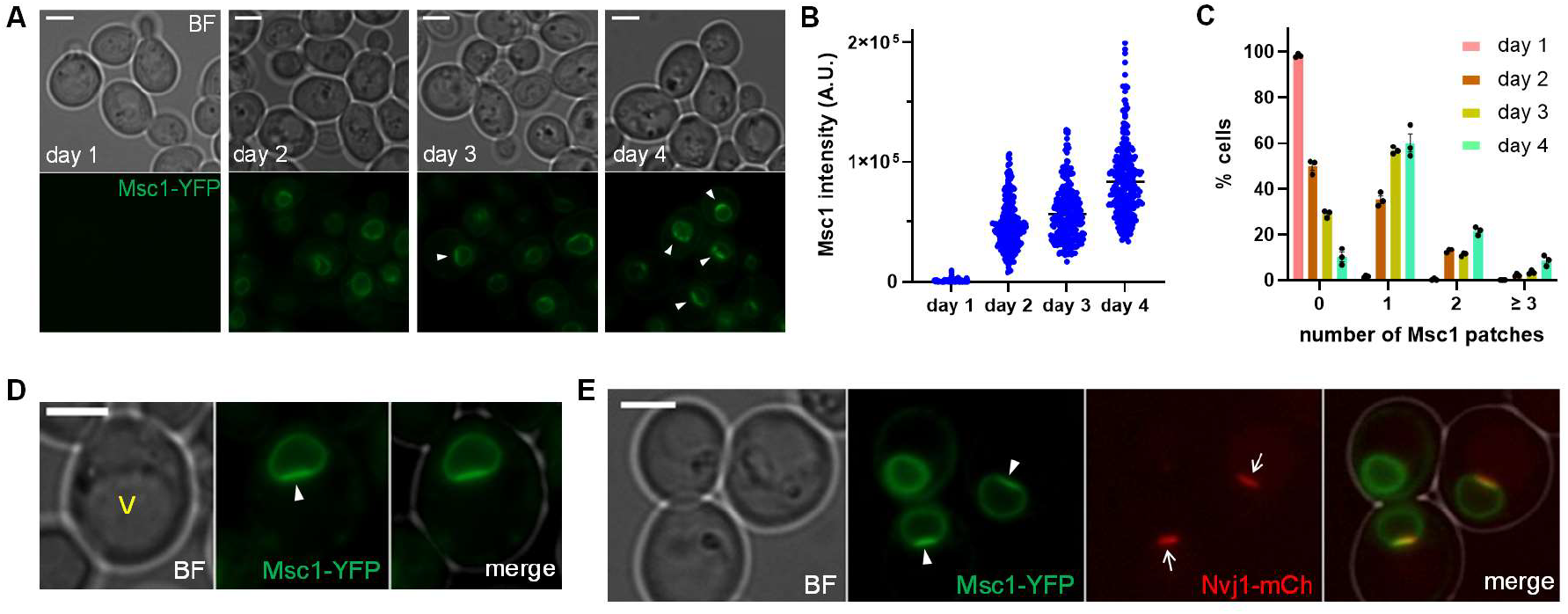
Msc1 patches increase after the log phase and colocalize with Nvj1 NVJs. (**A**) Micrographs of yeast cells expressing Msc1-YFP through four days of growth. (**B**) Single cell quantification of Msc1-YFP intensity at the nuclear envelope (NE) (pull of n=3). (**C**) Percentage cells with Msc1 NE patches (mean ± sem, n=3). (**D**) Micrograph of a Msc1 patch when the vacuole is visible through BF. Note how the patch apposes the vacuole surface. (**E**) Micrograph of three NE in cells that co-express Msc1-YFP and Nvj1-mCherry. Note the full co-localization of Msc1 patches and Nvj1, whereas Msc1 is evenly distributed in the absence of an NVJ. White arrowheads point to Msc1 patches. Scale bars correspond to 3 microns. BF, bright field; V, vacuole.

Remarkably, Msc1 patches often coincide with regions of the NE that dock with vacuoles, as seen through bright field (BF) microscopy (Figure 1D). The shape, location, and abundance of patches after passing the log phase led us to believe that they may colocalize with NVJs. Indeed, NVJs form after the diauxic shift, when inactivation of the nutrient-sensing kinases TORC1 and PKA coupled with the activation of Snf1 and Rim15 triggers the expression of the *NVJ1* gene (Galdieri et al., 2010; Gasch et al., 2000). In order to check whether Msc1 patches and NVJ colocalize, we constructed a strain co-expressing Msc1-YFP and Nvj1-mCherry. Nvj1 is the NE component of the NVJ. We observed a highly consistent colocalization of both proteins (Figure 1E). Indeed, in cells where Msc1 was already strong but Nvj1 was not present, the distribution of Msc1 was even, whereas Msc1 concentrated onto the Nvj1 patches when Nvj1 was present (90.72 ± 1.35 %; mean ± sem, n=3).

### Msc1 influences little on both the diauxic shift and post-diauxic growth

The timing of Msc1 enrichment suggests that Msc1 may play a role after the diauxic shift. In standard growth conditions in the nutrient-rich liquid YPD media under non-caloric restriction (2% glucose), the log phase lasts around one day and the post-diauxic phase up to seven days (Herman, 2002). To check whether Msc1 was important for the growth pattern after the log phase, we compared the growth curves of isogenic strains bearing a wild type copy of *MSC1* (WT) and its Δ*msc1* knockout counterpart. To fully assess all growth phases, including the diauxic shift and post-diauxic growth, we used a real-time mini biofermenter that reads optical density (OD) at 850 nm, which widens the linear range at higher cell densities. We observed that both strains grew rather similarly, including the apparent biomass at what the diauxic shift occurred (Figure 2A). In both strains, the log phase lasted ∼24h, with the post-diauxic growth spanning 2-3 days afterwards. The absence of Msc1 did impinge in this growth phase. There were, though, two slight differences between the strains; *msc1*Δ had both a longer lag phase at the beginning and appeared to yield a higher biomass in stationary phase proper (day 4 onwards) (Figure 2B and C). However, there were no more cells in the *msc1*Δ mutant than in the WT by the end of the experiment (1.8 vs. 2.07 × 10^8^ cells/mL, respectively).

**Figure 2.**
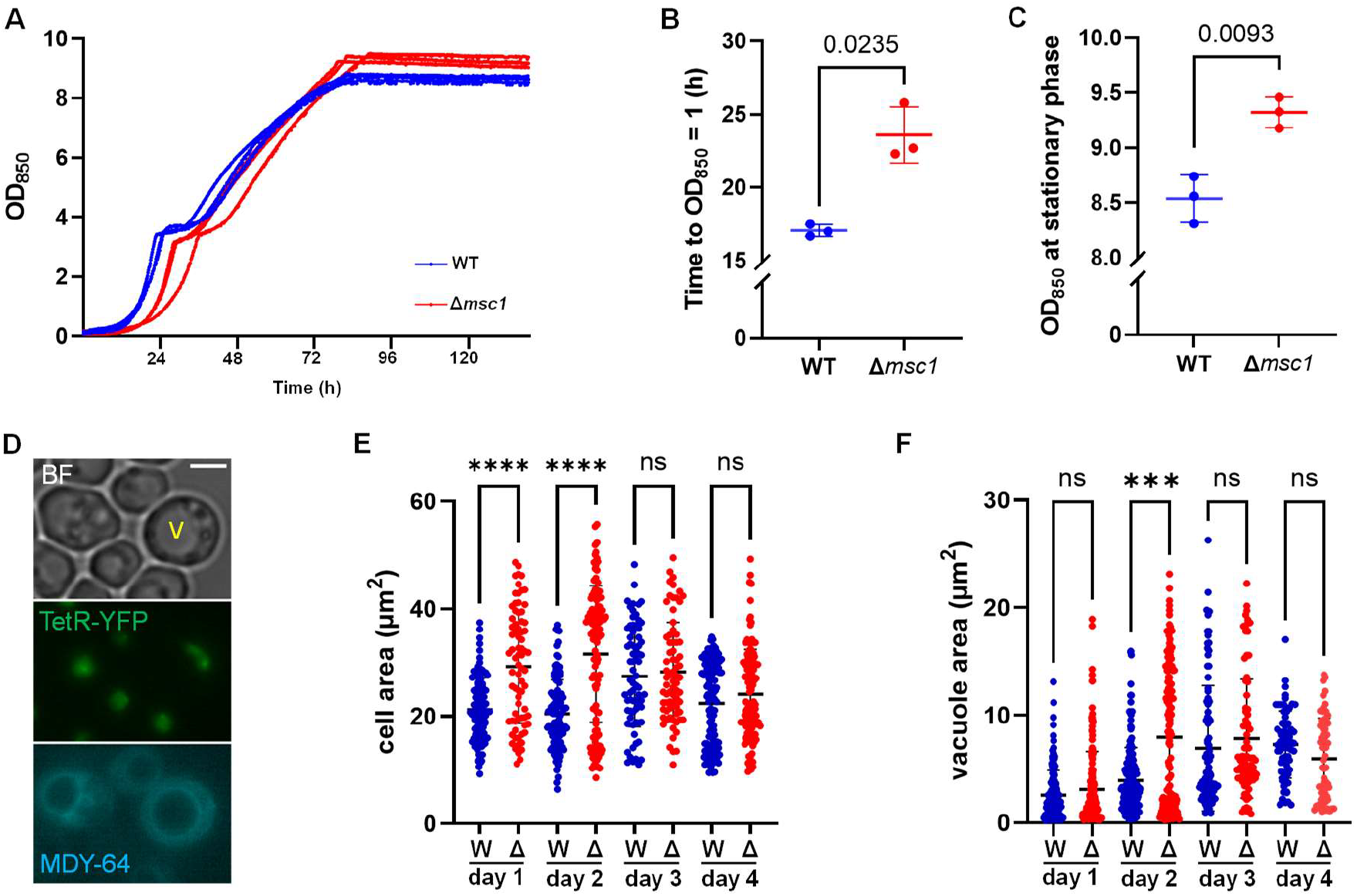
Growth and morphological cell profile in the absence of Msc1. **(A)** Comparative growth curves of the WT and Δ*msc1* strains (n=3). **(B)** Duration of the lag phase from the curves in A (mean ± sem, n=3). The time required to reach 1 OD_850_ was used as a proxy for the lag phase duration. (**C**) Maximum densities at the stationary phase (mean ± sem, n=3). (**D**) Representative micrograph of Δ*msc1* cells four days after inoculation (sum projection of six z slices). The strain labels the nucleoplasm (in green) and the right telomere of chromosome XII (as a stronger green spot) through the constitutive expression of the TetR-YFP and a *tetO* array located at the corresponding locus. MDY-64 labels the vacuolar membrane. Note the prominent vacuoles seen both by MDY-64 and in the bright field (BF) channel. Scale bars correspond to 3 microns. V, vacuole. (**E**) Single cell quantification of cell areas during growth. (**F**) Single cell quantification of vacuole areas. Note that the Δ*msc1* strain shows increased heterogeneity, especially in the first two days. For panels B and C, statistical comparisons were performed through the two-tailed unpaired Welch’s t test and actual p values are shown above. For panels E and F, statistical comparisons were performed through the Kruskal-Wallis test followed by Dunn’s multiple comparisons test (***, p<0.001; ****, p<0.0001).

After excluding an increase in cell number as the underlying cause of the higher OD_850_, we next took micrographs of cells along the growth curve, and measured both cell and vacuole area, since OD values depend not only on cell number but also on cell volume and density (refractive index) of cell content. To outline vacuoles, we used the endomembrane dye MDY-64, which tends to preferentially stain the vacuolar membrane (VM), yet it can also weakly stain the NE and the cell surface (endoplasmic reticulum and/or plasma membrane) (Matos-Perdomo et al., 2022). We found that WT cells showed more compact distributions in both cell and vacuole area than did *msc1*Δ cells during the first two days (Figure 2E and F). In general, Δ*msc1* populations displayed higher areas and a wider range relative to the WT, with a much clearer heterogeneity and non-gaussian subpopulation structure. This suggests that, even in the log and early post-diauxic phases, the Δ*msc1* strain exhibits cell distributions expected for stationary phase subpopulation. This is, cell size heterogeneity, mostly swollen unbudded cells with large vacuoles, likely as a result of vacuole fusion. As cells transit towards the end of the post-diauxic growth and the stationary phase proper (days 3 and 4), the differences between the strains became more subtle. Therefore, the higher OD_850_ value cannot be attributed to a larger cell volume and is probably due to changes in refractive index and/or density, which could be a consequence of changes in lipid composition, cell wall thickness or hyper-accumulation of storage glycogen granules or lipid droplets (Herman, 2002).

Finally, we determined the viability and vitality of the cells after 4 days of growth with propidium iodide and methylene blue, respectively, finding that more than 95% of the cells of both strains are still viable and metabolically active (Figures S1 and S2).

### Msc1 is important to promptly recover from quiescence

Next, we determined the colony-forming ability in a prolonged stationary phase through a clonogenic assay (Figure 3). For this purpose, cultures for both strains were incubated for a month in liquid YPD, and samples were taken through the days and compared for its ability to form colonies in fresh YPD plates. Since the stationary phase is characterized by a cell population that stops proliferation due to both exhaustion of essential nutrients and accumulation of inhibitory metabolic byproducts, and autophagy is mostly important to preserve viability under low nutrients, we grew the cultures in high aeration conditions that diminished the accumulation of toxic metabolites such as acetic acid (Dijken and Scheffers, 1986). Under agitated and well-oxygenated conditions, a greater proportion of the population enters quiescence (G0) and can remain viable for very long periods (up to months) without requiring changes to the medium (Gray et al., 2004). In general, *MSC1* and Δ*msc1* remained largely viable throughout (Figure 3A). The main difference occurred in the size of the new colonies (Figure 3B-D). The Δ*msc1* strain showed a steady decline in colony size after 14 days. By the end of the assay (28 days), the difference between the WT and *msc1*Δ was rather significant. This result indicated that Msc1 might be important for G0 cells to germinate promptly upon the return to favorable conditions, although its absence does not compromise overall viability in G0. This result is also consistent with our previous observation that *msc1*Δ spores germinate much slower than *MSC1* spores (Medina-Suárez and Machín, 2025).

**Figure 3.**
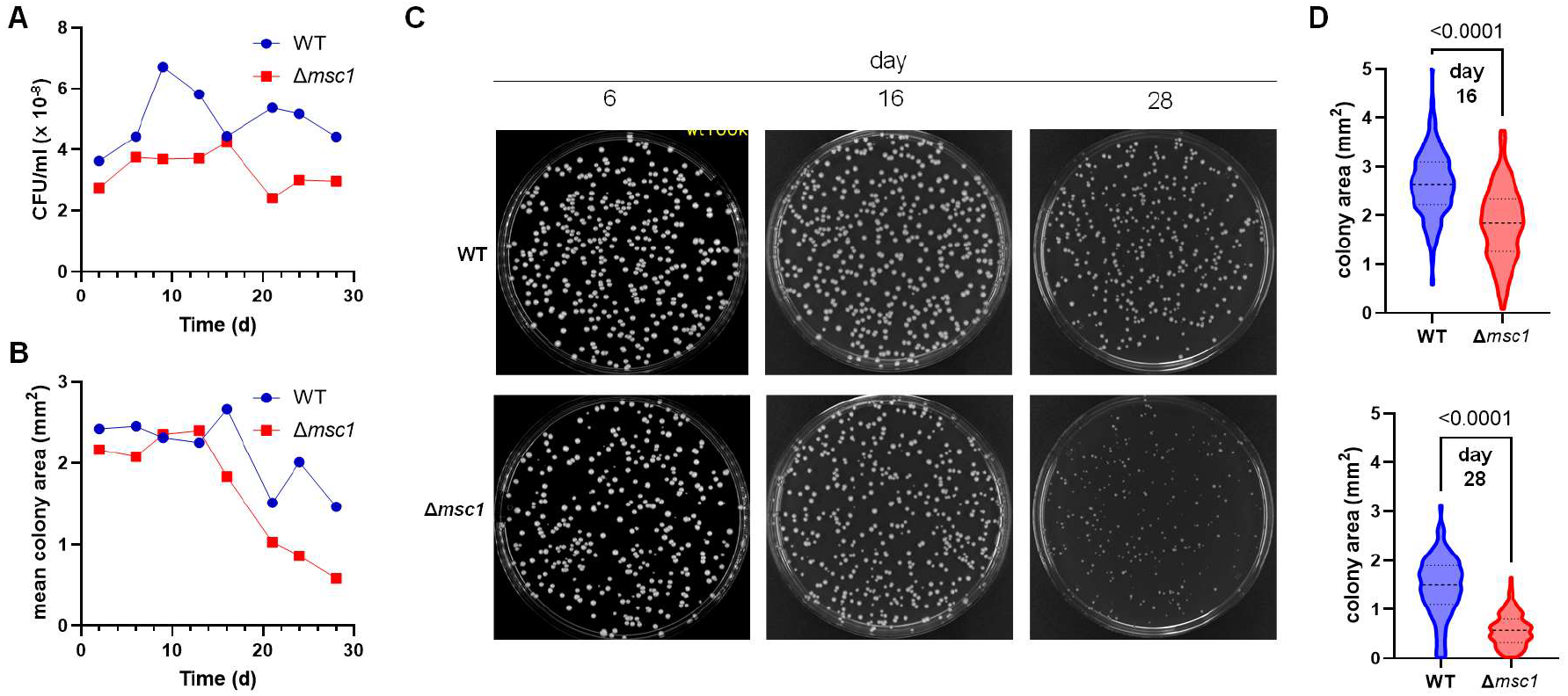
Fitness of Δ*msc1* in stationary phase. (**A**) Comparative colony-forming ability through the stationary phase as inferred from colony forming units (CFUs) (28 days of growth in YPD, it starts at day 2). (**B**) Comparative colony size of the same experiment. (**C**) Colonies after 4 days of growing in fresh YPD plates from samples taken at the indicated days of the experiment. (**D**) Quantification of colony areas of panel C (days 16 and 28). Statistical comparisons were performed through the two-tailed unpaired Welch’s t test and actual p values are shown above.

### Msc1 excludes nuclear pore complexes from nucleus-vacuole contact zones

A defining feature of the *msc1*Δ phenotype is the misdistribution of NPCs, which aggregate into both NE clusters resembling the Storage of Improperly assembled NPCs (SINC) and cytosolic foci (Medina-Suárez and Machín, 2025). To further characterize these Msc1-enriched NVJ subdomains, we investigated their spatial relationship with NPCs. While in cells co-expressing Msc1-YFP and Nup49-mCherry (a NPC Nup protein) both markers coincide when Msc1 is evenly distributed across the NE, we noticed a clear depletion of Nup49 in NE subdomains presenting Msc1 patches (Figure 4A). This pattern was observed in nearly all cases examined, and strongly suggests that there is an exclusion relationship between Msc1 and NPCs.

**Figure 4.**
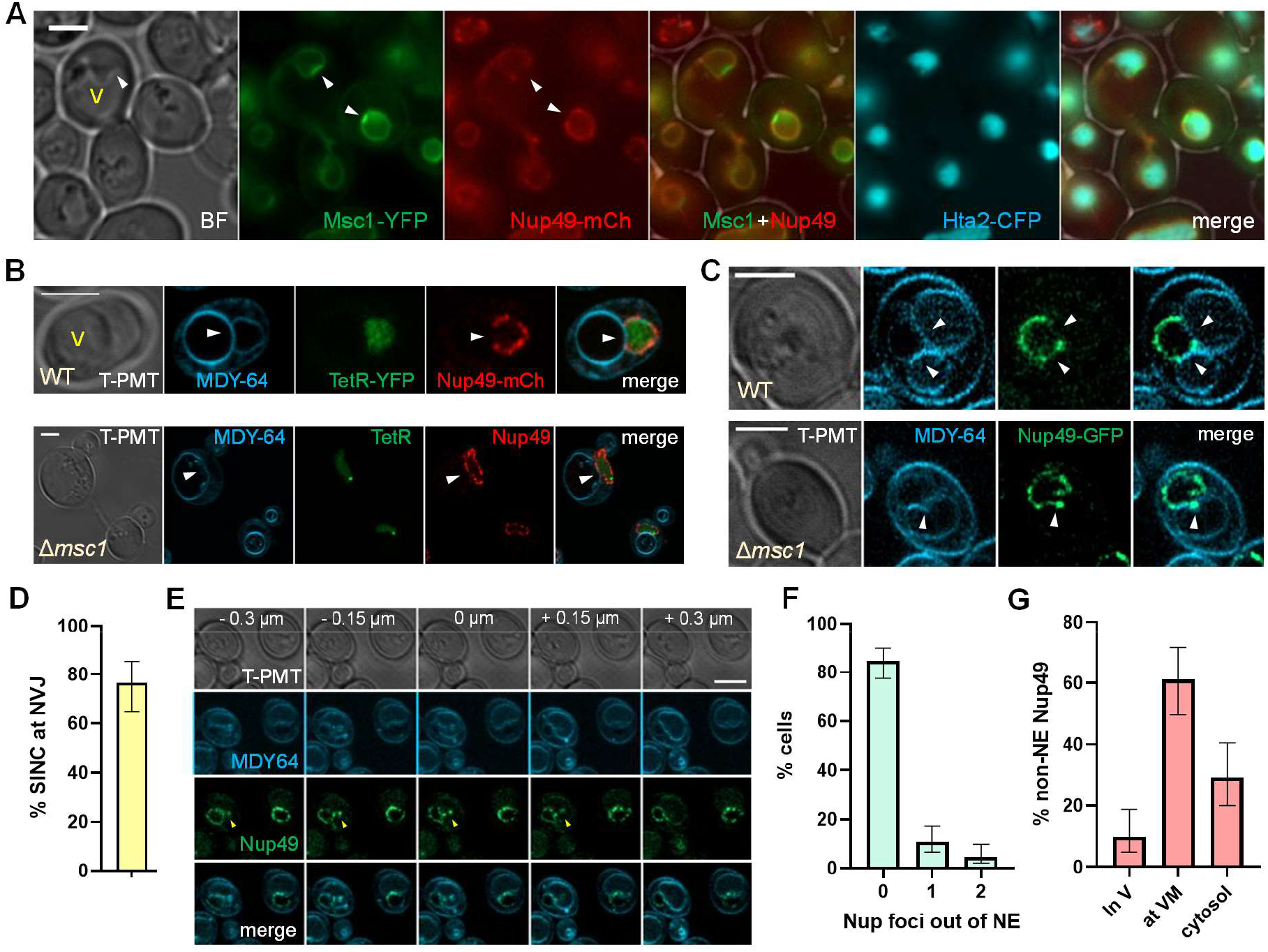
Msc1 excludes NPCs from nuclear envelope subdomains that contact vacuoles. (**A**) Micrographs of cells expressing Msc1-YFP, Nup49-mCherry (as a NPC marker) and Hta2-CFP (as a chromatin marker). Note that Nup49 is excluded from NE subdomains enriched in Msc1 (Msc1 patches). (**B**) Airyscan superresolution micrographs of representative post-diauxic cells of a wild type and a Δ*msc1* mutant that express Nup49-mCherry and TetR-YFP. Endomembranes were stained with the MDY-64 dye. Note how (i) the dye stains strongly the vacuolar membrane (VM) and less strongly the NE, and (ii) NPCs are present where VM and NE apposed in the Δ*msc1* strain. (**C**) As (B) but with strains that express Nup49-GFP. Note how in the Δ*msc1* example a Nup enriched foci (SINC) appears to invaginate into the vacuole. (**D**) Quantification of Nup49 SINCs located at NE-VM interacting zones in the Δ*msc1* strain (from the pool of two experiments, two days after inoculation; only cells with one SINC were counted, N= 64). (**E**) Walk through z slices of a cell that invaginates Nup49 into a large vacuole (yellow arrowheads). Slices taken with Airyscan 2 superresolution. (**F**) Percentage of cells with Nup49 aggregates out of the NE in the same experiments (N = 130 cells). (**G**) Subcellular distribution of Nup49 aggregates (N = 72 foci). Error bars represent CI95 of the proportions. White arrowheads point to NE-VM contact sites. Scale bars correspond to 3 microns. BF, bright field; V, vacuole; VM, vacuolar membrane.

To determine whether Msc1 was required for such an exclusion, we compared the presence and absence of Msc1 in two strains that express Nup49 tagged with different fluorescent proteins (mCherry and EGFP) along with nucleoplasm markers. In these strains, MDY-64 was again used to stain VMs and the NE. We confirmed that NPCs are excluded from NE-VM contact zones in the WT, but this was not the case for *msc1*Δ (Figure 4B and C). In *msc1*Δ, SINCs were also evident at the NE after the diauxic shift, whereas they were rare in WT cells. (Figure 4C, S3 and S4). A vast majority of these SINCs touch the vacuole (Figure S4A). The mean VM-NE contact perimeter was never higher than one third of the whole NE perimeter, but in cells where only one SINC was visible, 75% of SINCs locate at VM-NE contact zones (Figure 4D), which is far from what is expected by chance (P<0.0001, Fisher’s exact test, N=64). In addition, these SINCs often appeared to be associated with vacuolar invaginations (Figure 4C and S3), a morphology consistent with early PMN-like events. Even there were examples where these invaginations are large enough to pull NPCs far from the rest of the NE (Figure 4E).

Apart from the presence of SINCs, the absence of Msc1 also causes the formation of non-NE cytosolic Nup49 aggregates (Figure 4F). These aggregates are also found in very close proximity to the VM, suggesting that they may be a cargo for autophagy when cells have passed the log phase of growth (Figure 4G and S4).

Taken together, our new results support our previous observations on the role of Msc1 in surveilling the NE, potentially through the clearance of misassembled NPCs. In addition, our results strongly associate the enrichment of Msc1 at NVJs with the exclusion of NPCs. Thus, Msc1 might define territories that either host or exclude NPCs. This is in turn consistent with the role given to the Msc1 orthologue in *S. pombe* Les1, which excludes NPCs from the ends of the anaphase nucleoplasmic bridge (Dey et al., 2020). However, the alternative scenario of NVJs not being entirely functional in the absence of Msc1 cannot be disregarded, despite the observation of invaginating events of NPCs into the vacuole in *msc1*Δ. In this regard, a recent preprint points out that Msc1 might be important for fully functional NVJs (Mito et al., 2026). These observations raise the possibility that Msc1 might contribute to adaptive NVJ remodeling in order to maintain NPC homeostasis under starvation. Consistent with this idea, other authors have shown that NVJs expand and remodel under NPC assembly stress conditions to promote nuclear envelope homeostasis and vacuole-dependent degradation of specific nucleoporins (Lord and Wente, 2020).

## Materials and Methods

### Strains, growth, and experimental conditions

All yeast strains are derivatives of YPH499 (congenic with S288C). Strains and relevant genotypes are listed in Table S1. New strains were constructed through standard PCR-based procedures as reported before (Medina-Suárez and Machín, 2025).

Strains were routinely grown overnight in rich YPD medium (yeast extract 1% w/v, peptone 2% w/v, glucose 2% w/v) at 25°C with moderate orbital shaking (150 rpm; 25 mm orbit) in an air orbital incubator.

Growth curves were determined in two Biosan RTS-1C personal bioreactors. Wild type and mutant cultures were run in parallel, alternating strains between bioreactors across biological replicates. The bioreactor settings were 25 ºC, 1000 rpm, and change of spinning direction every 3 s. The culture volume was 10 mL and real-time growth was measured at OD 850 nm every 10 min. The initial inoculum was set at 0.01 OD 600 nm from a fresh overnight YPD preinoculum. Final cell concentration was determined with an hemocytometer.

Long-term survival during stationary phase was determined through a chronological aging-like experiment. A YPD overnight preinoculum was 1:100 diluted into 100 mL of fresh YPD, aliquoted into 50 mL Falcon-like tubes (10 mL/tube) and incubated tilted for one month. At the indicated days (starting ∼48 h after inoculum), one tube was taken, the cultures were diluted 1:10,000 and 1:100,000 times, and 100 μL plated on YPD agar plates. These plates were further incubated for four days before taking photographs. Colony number and size were determined with ImageJ as follows: An intensity threshold was applied to identify colonies as objects, and the watershed function was used to separate juxtaposed colonies. Proper separation was checked visually and corrected when required.

### Bright field and epifluorescence microscopy

Two inverted epifluorescence microscopes were used as reported before (Matos-Perdomo et al., 2022): A Leica DMI6000B with a Xenon excitation lamp, a DFC350 digital camera, and a 63X/1.30 immersion objective; and a Zeiss Axio Observer.Z1/7 equipped with Colibri-7 LED excitation system, an Axiocam 702 sCMOS camera, and a 63X/1.40 immersion objective. In both cases, narrow-band filter cubes were used for co-visualization of CFP, YFP/GFP, and mCherry with minimal emission crosstalk.

Whenever possible, cells were imaged alive or shortly after staining with dyes. Briefly, 200 μl of cell culture was collected at each time point, centrifuged at 300 x g for 1 min at room temperature, the supernatant carefully retired, and ∼1.5 μl of the pellet was placed on the microscope slide. For each field, we captured a series of 6–14 z-focal plane images (0.2–0.4 μm spacing between each consecutive planes), and then we processed images with the Leica AF6000, Zeiss Zen Blue and ImageJ software. For z-stack 2D projections, we usually used the maximum intensity method.

When endomembrane staining was required, a 200 μl sample of the culture was pelleted at 500 x g for 1 min, and 1 μl of the pellet was mixed directly on the slide with 0.5 μl of a 3X MDY-64 solution (10 μM final concentration). Alternatively, 8 μl of a 25X solution was added to 200 μl of the culture, incubated 15 min, washed twice with either phosphate buffer saline (PBS) or 10 mM Hepes pH 8, and pelleted. To assess plasma membrane integrity, cells were incubated with 3 μg/mL of PI for 10 min, washed with PBS twice and directly visualized on the red channel. For metabolic activity, cells were stained with 0.04% w/v methylene blue for 10 min, washed with PBS and visualized in a bright field Leica DM2000 LED microscope equipped with a Flexacam i5 color camera.

### Confocal super-resolution microscopy

A Zeiss LSM980 equipped with Airyscan 2 was used for super-resolution microscopy as reported before (Matos-Perdomo et al., 2022). Briefly, samples were gathered as above, and we used the following laser lines for excitation of fluorescent tags: 405 nm for CFP and MDY-64; 514 nm for YFP; 488 nm for GFP; and 561 nm for mCherry. Bright-field (BF) images were acquired with a T-PMT detector (pinhole 1 AU). After imaging, Airyscan processing was conducted.

Approximately 30 z planes (with 0.15 μm spacing) were obtained across the entire field.

### Data processing and statistics

Quantification of images was performed with Fiji/ImageJ (NIH). Msc1 intensities at the NE were determined as previously described (Medina-Suárez et al., 2024). Phenotype categorization and counting were performed manually. In all cases, more than 100 cells were counted per condition and experiment.

Error bars indicate the standard error of the mean (SEM), standard deviation (SD), or 95% confidence intervals (95% CI), as specified in each figure legend. Sample sizes are reported as independent biological replicates (n) or analyzed events/cells (N). GraphPad Prism 10 was used for graph generation and statistical analyses. Statistical tests are indicated in the corresponding figure legends.

## Acknowledgements

We thank Samantha Morais-Armas for technical help and discussion.

## Author contributions

Sara Medina-Suárez: Formal analysis, Investigation, Methodology, Validation, Visualization, Writing–review and editing.

Héctor M Estévez-Silva: Investigation, Methodology, Validation, Writing–review and editing. Noelia Rodríguez-Herrera: Formal analysis, Visualization, Writing–review and editing.

Félix Machín: Conceptualization, Data curation, Formal analysis, Funding acquisition, Investigation, Methodology, Project administration, Resources, Supervision, Validation, Visualization, Writing–original draft, Writing–review and editing.

## Data availability

All data is contained within the manuscript. Raw data and materials are available from the corresponding author upon reasonable request.

---

**Figure S1.**
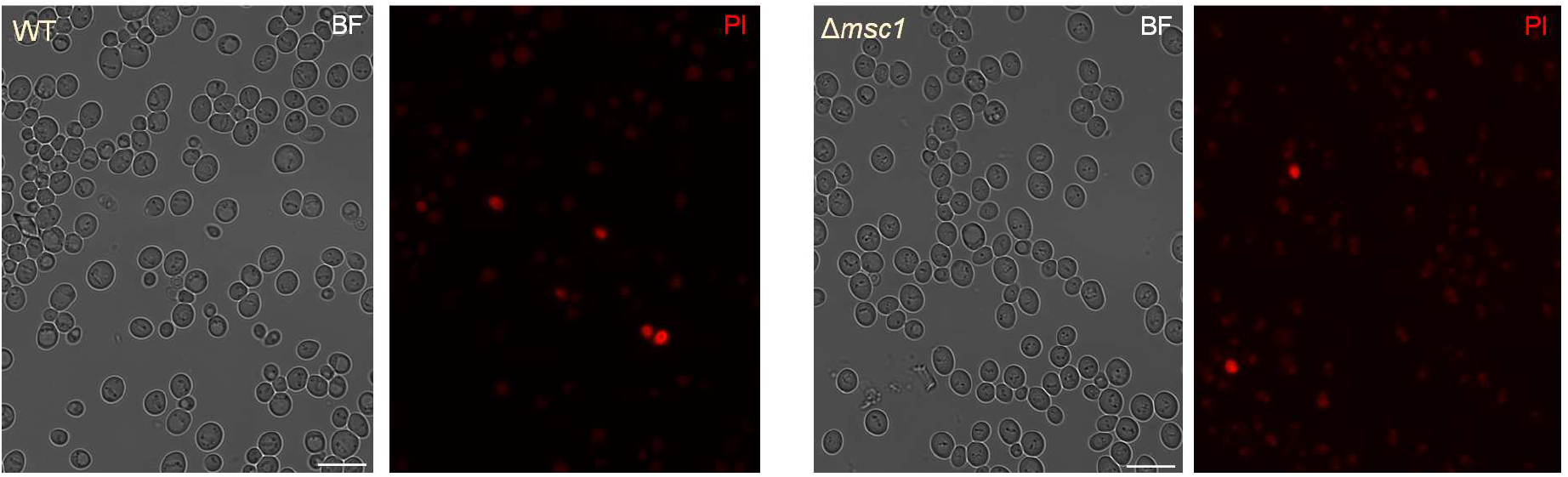
Cells remain viable in early stationary phase in the absence of Msc1. Related to Figure 2. Samples in the fourth day of growth in YPD were stained with propodium iodide (PI) and visualized. Note that more than 95% of cells do not stain for PI.

**Figure S2.**
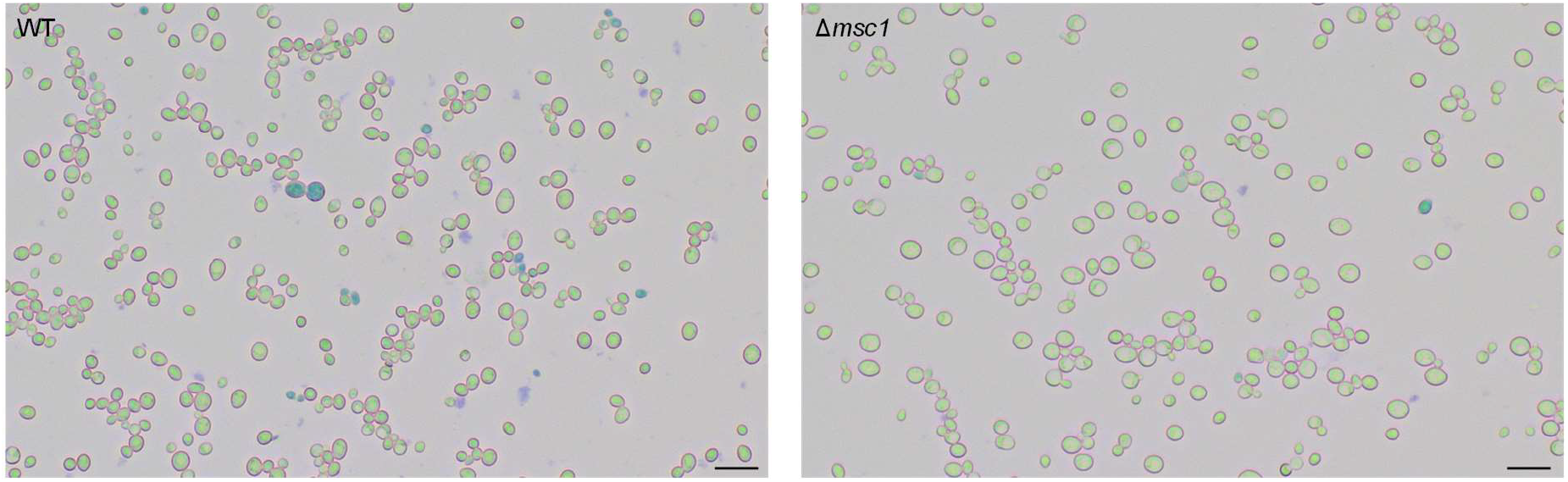
Cells vitality remains high in early stationary phase in the absence of Msc1. Related to Figure 2. Samples in the fourth day of growth in YPD were stained with methylene blue and visualized in the bright field. Note that more than 95% of cells do not turn into blue.

**Figure S3.**
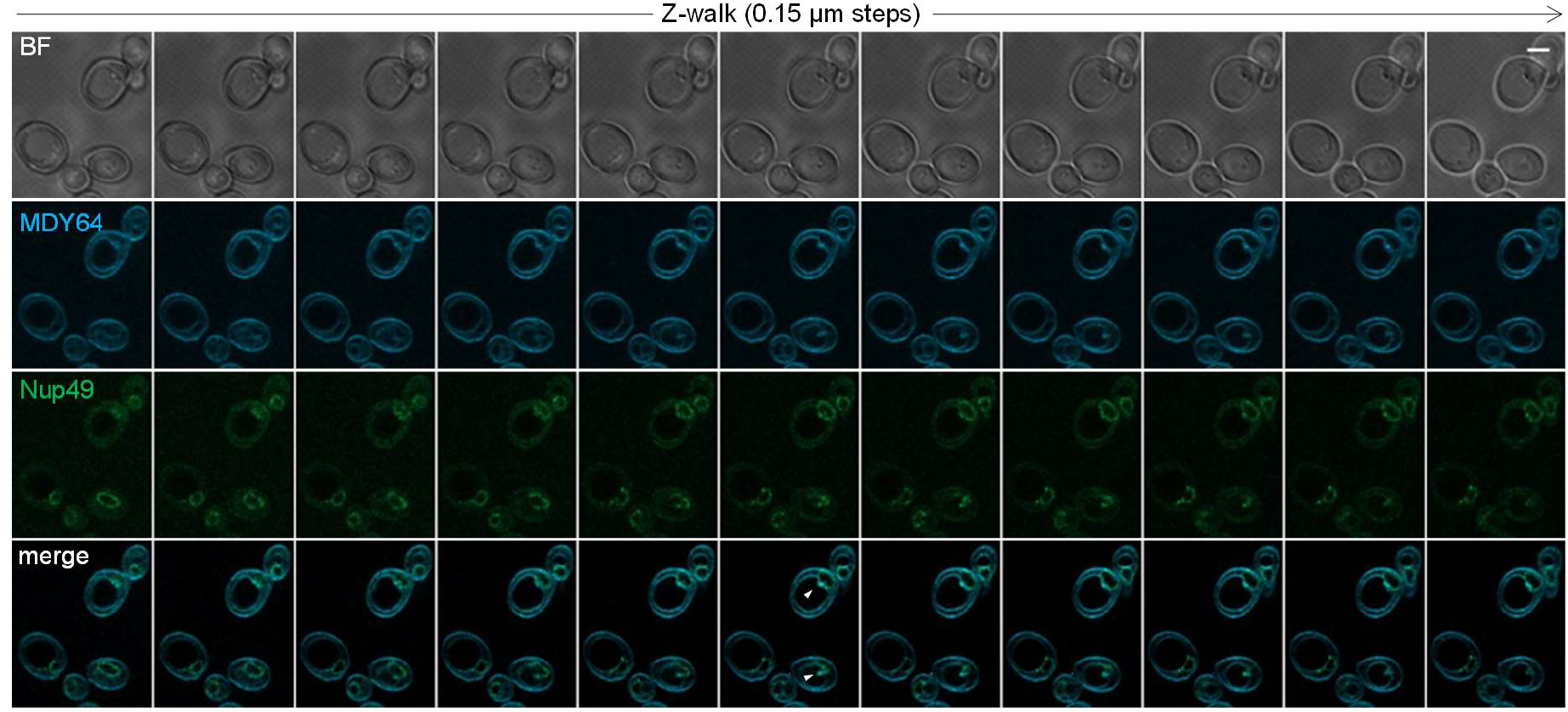
NPCs invaginate into vacuoles in the absence of Msc1. Related to Figure 4. Walk through z slices of two cells that start invaginating SINC into vacuoles (arrowheads). Slices taken with Airyscan 2 superresolution. Scale bars correspond to 3 microns. BF, bright field (through a T-PMT detector).

**Figure S4.**
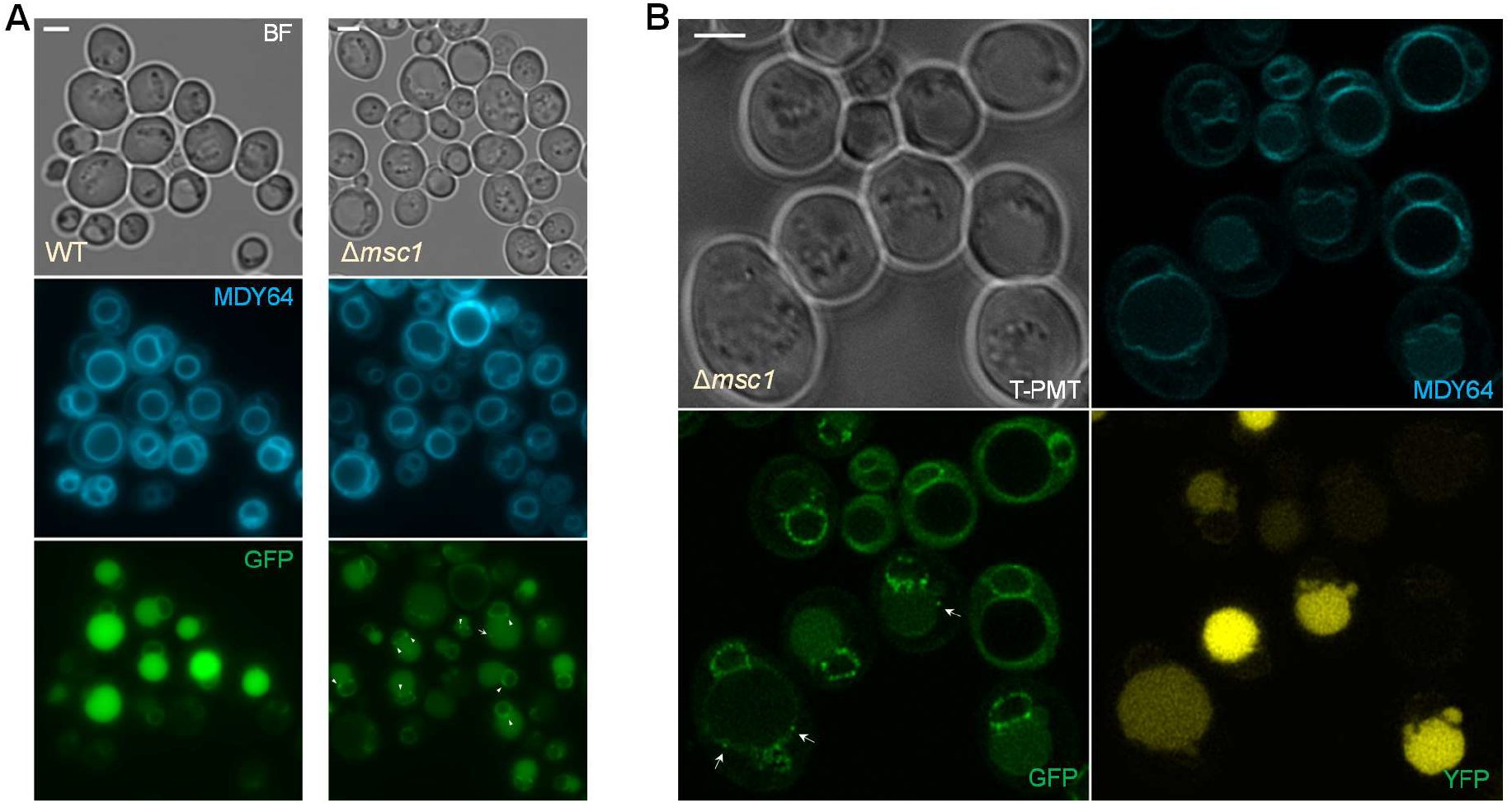
NPC SINCs and cytosolic Nup aggregates accumulate in early stationary phase in the absence of Msc1. Related to Figure 4. WT and Δ*msc1* strains expressing Nup49-GFP were grown for 4 days before taking micrographs. (**A**) Epifluorescence images of both strains. (**B**) Airyscan 2 superresolution of Δ*msc1*. Note that these strains develop a strong vacuolar autofluorescence as cells approach quiescence. White arrowheads point to SINC. Arrows point to extranuclear Nup aggregates. Scale bars correspond to 3 microns. BF, bright field.

**Table S1.** Strains used in this work.

| Name | Genotype<sup>1</sup> |
| --- | --- |
| FM2684 | *MATa bar1Δ leu2-3,112 ura3-52 his3-Δ200 trp1-Δ63 ade2-1 lys2-801; cdc15-2:9myc::HphNT1; Msc1:eYFP::KanMX* |
| FM2070 | *MATa bar1Δ leu2-3,112 ura3-52 his3-Δ200 trp1-Δ63 ade2-1 lys2-801; ade2-1::TetR:YFP::ADE2; tetO(5.6Kb)::1061Kb-ChrXII::HIS3; cdc15-2:9myc::HphNT1* |
| FM2728 | FM2070; *NUP49:mCherry::kanMX* |
| FM2807 | FM2070; *Δmsc1::NatNT2* |
| FM2875 | FM2728; *Δmsc1::NatNT2* |
| FM2876 | *MATa bar1Δ leu2-3,112 ura3-52 his3-Δ200 trp1-Δ63 ade2-1 lys2-801; cdc15-2:9myc::HphNT1; Msc1:eYFP::KanMX; Nvj1-mCherry::NatNT2* |
| FM2947 | *MATa bar1Δ leu2-3,112 ura3-52 his3-Δ200 trp1-Δ63 ade2-1 lys2-801; cdc15-2:9myc::HphNT1; NUP49:eGFP::TRP1* |
| FM2954 | FM2947; *Δmsc1::NatNT2* |
| FM3268 | *MATa bar1Δ leu2-3,112 ura3-52 his3-Δ200 trp1-Δ63 ade2-1 lys2-801; cdc15-2:9myc::KanMX; Nup49-mCherry::NatNT2; Hta2:eCFP::TRP1, Msc1:eYFP::Hph* |

<sup>1</sup> Only relevant genotypes for this work are indicated. A non-italic beginning refers to a parental strain. Semicolon (“;”) separates genetic modifications accomplished through transformation.

